# Inhibition of inositol-requiring enzyme 1α RNase activity protects pancreatic beta cell and improves diabetic condition in insulin mutation-induced diabetes

**DOI:** 10.1101/2020.08.30.274381

**Authors:** Oana Herlea-Pana, Venkateswararao Eeda, Ram Babu Undi, Iulia Rus, Hui-Ying Lim, Weidong Wang

## Abstract

Proinsulin misfolding in the endoplasmic reticulum (ER) plays an important role in β-cell dysfunction and death and the pathogenesis of mutant *INS*-gene-induced diabetes of youth (MIDY). There is no effective treatment for MIDY except the insulin administration. Here, we found that the ER stress sensor inositol-requiring enzyme 1α (IRE1α) was activated in the Akita mice, a mouse model of MIDY. Normalization of IRE1α RNase hyperactivity by pharmacological inhibitors significantly ameliorated the hyperglycemic conditions and increased serum insulin levels in Akita mice. These benefits were accompanied by a concomitant protection of functional β-cell mass, as shown by the suppression of β-cell apoptosis, increase in mature insulin production and reduction of proinsulin level. At the molecular level, we observed that the expression of genes associated with β-cell identity and function was significantly up-regulated and ER stress and its associated inflammation and oxidative stress were suppressed in islets from Akita mice treated with IRE1α RNase inhibitors. This study provides the first evidence of the in vivo efficacy of IRE1α RNase inhibition in Akita mice, pointing to the possibility of targeting IRE1α RNase as a therapeutic direction for the treatment of MIDY diabetes.

## INTRODUCTION

In pancreatic β-cells, proinsulin, the precursor to insulin, is folded in the endoplasmic reticulum (ER) and trafficked to the Golgi for processing into mature insulin during biosynthesis. However, proinsulin is misfolding-prone; even under normal physiologic condition, up to 20% of newly synthesized proinsulin fail to fold properly (1-4). The portion of misfolded proinsulin increases further under conditions of insulin mutations (5, 6), high demand on insulin production (e.g., obesity-linked insulin resistance) (7-10) or perturbed ER folding environment (e.g., hyperglycemia, lipotoxicity, oxidative stress) (2, 3, 11). In the case of insulin mutation, a number of single point mutations in the insulin gene have been identified in patients of Mutant *INS*-gene-induced Diabetes of Youth (MIDY) who exhibit varying degrees of diabetes symptoms (6, 12). A majority of these mutations cause the misfolding of mutant proinsulin in the ER of β-cells, which not only interferes with the folding of wild-type proinsulin but also induces ER stress, a condition of accumulation of unfolded or misfolded proteins in the ER, progressively leading to β-cell dysfunction and death (4-6, 12, 13). Notably, proinsulin misfolding is regarded as one of the primary initiating factors of ER stress in β-cells (7, 14, 15).

Upon ER stress, the unfolded protein response (UPR) is initially activated to serve as an adaptive means to resolve ER stress, but will eventually become maladaptive when activated chronically. The UPR is transduced by three core pathways – inositol requiring enzyme 1-α (IRE1-α), activating transcription factor 6 (ATF6), and PKR-like ER kinase (PERK). IRE-1α, the most evolutionarily conserved among the UPR sensors, is an ER transmembrane protein with dual serine/threonine kinase and RNase domains. When misfolded proteins bind to its luminal domain, IRE1α monomers aggregate in the ER membrane, juxtaposing cytosolic kinase domains and facilitating kinase trans-autophosphorylation (16-19) and enabling its RNase activity. Activated IRE1α RNase unconventionally splices XBP-1 mRNA and initiates a process termed regulated IRE1-dependent decay of RNA (RIDD), in which select RNAs are degraded that eventually causes cell death. IRE1α hyperactivation has been observed to contribute to pathological manifestation and progression (20-23), and overexpression of IRE1α alone is sufficient to cause cell death (24, 25). As ER stress has been detected in multiple cell types including β-cells that are important in the pathogenesis of diabetes, targeting IRE1α or ER stress has been proposed as an important therapeutic direction for diabetes (19, 26).

On the other hand, IRE1α is known to play important roles in the maintenance of ER homeostasis in physiological settings as well as in the early adaptive phase of ER stress (20, 27-32). The IRE1α/XBP1 axis is critical for ER expansion in secretory cells such as plasma cells (33). Moreover, IRE1α prevents ER membrane permeabilization and ER stress-induced cell death under pathological conditions (28). IRE1α also plays an important role in the regulation of postprandial biosynthesis, proinsulin folding, and secretion of insulin and protection in β-cells (27, 32, 34, 35); as a result, IRE1α knockout β-cells exhibited functional impairment (35, 36). In addition, in some settings, IRE1α activation is known to subside in cells under chronic exposure to synthetic ER stressors in vitro. For instance, in the liver of high fat diet-fed mice IRE1α RNase activity is down-regulated (37, 38). Together, these findings support an important physiological role of IRE1α and raise the question as to whether inhibiting IRE1α represents a viable approach in countering ER stress-related pathological diseases.

In this study, we report, for the first time, the effect of IRE1α RNase inhibition on the diabetic conditions of the *Akita* mouse (39-41), an animal model of MIDY. We showed that treating *Akita* mice with IRE1α RNase inhibitors [STF-083010 (STF) and 4μ8C] significantly lowered blood glucose levels while increasing serum insulin levels. These beneficial effects were accompanied by a concomitant preservation of functional β-cells and suppression of β-cell death. Finally, ER stress markers, in particular, genes associated with IRE1α activity were suppressed in STF-treated β-cells. Collectively, these studies indicate that inhibition of IRE1α significantly improved the β-cell dysfunction and death, insulin deficiency and hyperglycemia observed in Akita mice, which could serve as a foundation for targeting IRE1α as a therapeutic direction for the treatment of diabetes.

## RESEARCH DESIGN AND METHODS

### Animal studies

C57BL/6J wild-type (WT) mice and Akita mice were obtained from Jackson laboratory (Bar Harbor, ME). The genotyping of Akita mice was confirmed using tetra-primer ARMS-PCR approach (41). Mice were housed in cages on a 12 h light (6:00 a.m. to 6:00 p.m.)−12 h dark (6:00 p.m. to 6:00 a.m.) cycle at an ambient temperature of 22 °C. All animals had access to normal chow diet and water ad libitum. All procedures involving animals were performed in accordance with the protocol approved by the Institutional Animal Care and Use Committee of the University of Oklahoma Health Science Center. All experiments were performed with age-matched female mice.

For Akita model, mice at 5-6 weeks of age were injected i.p. with either vehicle (*n* = 9 mice), STF (10 mg/kg body weight; 2 mg/ml in 10% DMSO in saline buffer; *n* = 9 mice) or 4μ8C (10 mg/kg of body weight) once daily. Compounds were dosed approximately 3−4 h before the initiation of the dark cycle (2−3 p.m. during the light cycle). Age-matched WT mice with no treatment serve as controls. Blood glucose levels were measured using the OneTouch Ultra2 glucometer after animals were fasted for 6 h and blood was obtained by tail snip. Measurements of body weight were performed weekly. At the end of the treatment period, mice were fasted for 4 h and euthanized, and pancreata were removed and weighted; for half the pancreata, a small portion of the tail end was cut and saved for insulin and proinsulin content measure while the remaining parts of these pancreata and the remaining whole pancreata were directly fixed in formalin, and paraffin-embedded.

### Glucose tolerance test and insulin tolerance test

Intraperitoneal glucose tolerance test (ipGTT) were performed after 16-h overnight fasting. Blood glucose levels were measured at 0, 15, 30, 60, and 120 minutes after an intraperitoneal administration of glucose at dose of 1.5 g/kg body weight. Intraperitoneal insulin tolerance test (ipITT) were performed after 4 hour fasting. Blood glucose levels were measured at 0, 15, 30, 60, and 120 minutes after an intraperitoneal administration of insulin at dose of 0.75 IU/kg body weight.

### Islet isolation procedure

Islets were isolated using the standard collagenase digestion method. Briefly, the pancreatic duct was cannulated and distended with Collagenase P (0.5 mg/ml, Sigma-Aldrich, USA) in 1x Hank’s balanced salt solution. Pancreata were incubated in water bath at 37C for 25 m. The reaction was stopped using RPMI 1640 with 10% fetal bovine serum. Islets were separated from the exocrine tissue using Histopaque-1077 (Sigma-Aldrich, USA). Hand-picked islets were cultured overnight at 37C in RPMI-1640 media containing 10% FBS (Fisher Scientific) before use in experiments.

### RNA isolation and qRT-PCR

Total RNA was extracted using TRIzol reagent (Invitrogen, Carlsbad, CA) according to the manufacturer’s protocol, and 2 μg of total RNA was reverse transcribed using a Superscript kit (Invitrogen). Regular RT-PCR was performed and the products was resolved by agarose gel electrophoresis. The full-length (unspliced, XBP1u) and spliced (XBP1s) forms of XBP1 mRNA were quantified using ImageJ software (National Institutes of Health, Bethesda, MD). Real-time PCR was performed in 96-well format using SYBR Select Master Mix (Applied Biosystems, Foster City, CA) with a CFX96 Real-Time PCR detection system (Bio-Rad, Hercules, CA). The amplification program was as follows: initial denaturation at 95°C for 15 min, followed by 45 cycles of 95°C for 15 s, 60°C for 1 min, and 40°C for 30 s. Relative mRNA levels were normalized against the housekeeping gene Cyclophilin A using the the ΔΔCt method. The primer sequences used were included as supplemental material in supplemental table 1.

### Glucose-stimulated insulin or proinsulin secretion

20 primary islets isolated from Akita mice treated with STF or vehicle were seeded in 96-well plates overnight. Islets were then incubated in fresh KRBH buffer (115 mM NaCl, 5 mM KCl, 24 mM NaHCO_3_, 2.5 mM CaCl_2_, 1 mM MgCl_2_, 10 mM HEPES, 2% w/v BSA, pH 7.4) containing 2.5 mM glucose for 1 h. Cells were incubated for an additional hour in KRBH buffer containing 2.5 or 16.7 mM glucose. The secreted insulin and proinsulin from supernatant were measured with insulin ELISA kits (ALPCO, Salem, NH) and Rat/Mouse Proinsulin kit (Mercodia), respectively. islets were lysed with RIPA buffer (50 mM Tris HCl pH 7.4, 1% NP-40, 0.25% sodium deoxycholate, 150 mM NaCl), and total cellular protein was determined with a Bradford protein assay. The secreted insulin levels were normalized for total protein.

### Insulin and proinsulin content measurements

For insulin or proinsulin content measurement, islets or pancreatic tissues was incubated and homogenized in 1.5% HCl in 70% EtOH overnight at -20C, and the solution was neutralized with equal volume of 1M Tris pH 7.5. Insulin and proinsulin were measured by ELISA using Mouse Insulin (ALPCO) and Rat/Mouse Proinsulin kit (Mercodia), respectively, and normalized to weights of pancreas for pancreatic tissues or to protein levels for islets.

### Immunofluorescent staining and islet mass measurement

At the end of the treatment period, mice were sacrificed and the pancreases were removed, fixed in formalin, and paraffin-embedded. For islet mass measurements, at least 3 mice from each group were analyzed. 6-8 slide sections on average from each mouse were sectioned with the separation at 150 μm increments. Sections were stained with an anti-insulin antibody (A0564, 1:500; Dako), anti-glucagon antibody (G2654, 1:500; Sigma), and DAPI (0.5 μg/mL). Alexa Fluor 488-, 555-, and 647-conjugated secondary antibodies (Jackson ImmunoResearch) were used. Images covering the entire tissue sample were subsequently captured in each section. The entire pancreas tissue, glucagon^+^, and insulin^+^ areas in each image were measured using ImageJ software (National Institutes of Health, Bethesda, MD). Relative β-cell area was defined as the sum of the islet β-cell area measured in all the images from each mouse divided by the sum of the total pancreas tissue area measured in all the images from each mouse, and was normalized by designating the β-cell area of wild-type B/6 mice as 1. All images were taken with an Olympus FV1000 confocal microscope and quantified with Image-J histogram software. Mouse anti-caspase 3 (cat# 9446, 1:500, CST), rabbit anti-Ki67 (Ab15580, 1:250, Abcam), mouse anti-proinsulin (GS-9A8, 1:100, Developmental Studies Hybridoma Bank), or Anti-4-Hydroxynonenal antibody [HNEJ-2] (ab48506, Abcam) was co-immunostained with anti-insulin antibody (A0564) and/or anti-glucagon antibody (G2654) in pancreatic sections.

### TUNEL staining

TUNEL (Terminal deoxynucleotidyl transferase-mediated nick-end labeling) staining was performed in pancreatic sections with In Situ Cell Death Detection Kit-Fluorescein (Roche) according to the manufacturer’s instructions. Anti-insulin antibody (A0564) and DAPI were used for β-cell marker and nuclear counter-staining. At least 50 islets for each of three mice were counted. All images were taken with an Olympus FV1000 confocal microscope and quantified with Image-J histogram software.

### Transmission Electron microscopy (TEM)

Mouse islets were isolated from animals as described in Animal studies section. Mouse islets were fixed with 0.1 M sodium phosphate buffer (pH 7.2) containing 2 % glutaraldehyde and 2 % paraformaldehyde for 1 h, then exposed to 2 % osmium tetroxide, stained with 2 % uranyl acetate, dehydrated with ethanol, and embedded in Epon (TAAB). Ultra-thin sections were stained with uranyl acetate and lead citrate, and images were recorded with a Hitachi H-7600 transmission electron microscope (Hitachi).

### Statistical Analysis

All values are reported as mean ± SEM. Statistical significance between groups was analyzed by unpaired Student *t* test or one-way ANOVA with a least significant difference test for multiple comparisons. The difference between groups was considered statistically significant if a *P* value <0.05.

## RESULTS

### Up-regulation of IRE1α RNase activity in pancreatic islets in Akita mice

The C96Y missense mutation in the Ins2 gene in Akita mice causes mutant proinsulin protein misfolding that is responsible for ER stress (4). Previously, the ER stress response markers PERK and ATF6 have been reported to be up-regulated in in vitro cultured immortalized β-cell lines carrying the Ins2^Akita/+^ mutation and islets freshly isolated from Akita mice (39, 42, 43). Consistent with this, we observed that the mRNA levels of the PERK pathway genes *ATF4* and *CHOP* and the ATF6 target gene *Bip* were up-regulated in islets freshly isolated from Akita mice over the age of 3 weeks (Fig. S1A-S1C). However, how IRE-1α responds to this mutation in β-cells in vivo is unclear. Earlier studies using in vitro β-cell lines yielded controversial results; in one report, using immortalized β-cell lines carrying Ins2^Akita/+^ mutation, the IRE-1α-XBP1 pathway was shown to be activated in Ins2^Akita/+^ β-cell lines compared to β-cell lines carrying wild-type Ins2 gene (44), whereas others reported that IRE-1α activity was down-regulated in stable β-cell lines expressing Ins2^Akita/+^ mutation (45). To investigate the in vivo IRE-1α activity in Akita islets, we first examined the splicing of *Xbp1* mRNA, a direct target of IRE-1α RNase. RT-PCR amplification of the *XBP1* mRNA followed by electrophoretic separation of spliced (*XBP1-s*) and unspliced (*XBP1-u*) forms of *XBP1* revealed a gradual increase in *Xbp1* mRNA splicing in the islets isolated from Akita mice from age of 2 weeks onwards compared to the age-matched WT mice (Fig. 1A). Quantitation of the ratio of *XBP1-s* to total *XBP1* (XBP1-u + XBP1-s) mRNA levels further corroborated the notion of increased Xbp1 mRNA splicing (Fig. 1A’). Quantitative RT-PCR (qRT-PCR) using a pair of XBP1 splicing-specific primers further showed that the spliced *Xbp1* mRNA levels were significantly increased in the Akita islets over a period of 12 weeks while total *Xbp1* mRNA levels increased only slightly during the same period (Fig. 1B, 1C). Second, as XBP1 protein translated from the spliced *XBP1* mRNA controls the transcription of a number of downstream genes that are involved in protein folding, we investigated changes in the transcript levels of the XBP1 target genes *EDEM1* and *P58*, in Akita islets. We observed that the mRNA levels of *EDEM1, P58*, and *ERDj4* were also markedly up-regulated in Akita islets over the age-matched counterparts (Fig. 1D, 1E), as assessed by qRT-PCR. Third, as hyperactivation of IRE-1α was proposed to lead to the initiation of IRE1-dependent decay of mRNA (RIDD), we analyzed the mRNA levels of *Blos1* and *Col6a1*, two typical RIDD target mRNAs by qRT-PCR. We observed that *Blos1* and *Col6a1* mRNA levels decreased progressively in the islets of Akita mice from 3-week old onwards (Fig. 1F, 1G). Together, our results demonstrate that IRE1α activity was already elevated at around 2 weeks of age, prior to the development of hyperglycemia in Akita mice, and that further elevation of its activity continued until the Akita mice developed overt diabetes.

**Figure 1.**
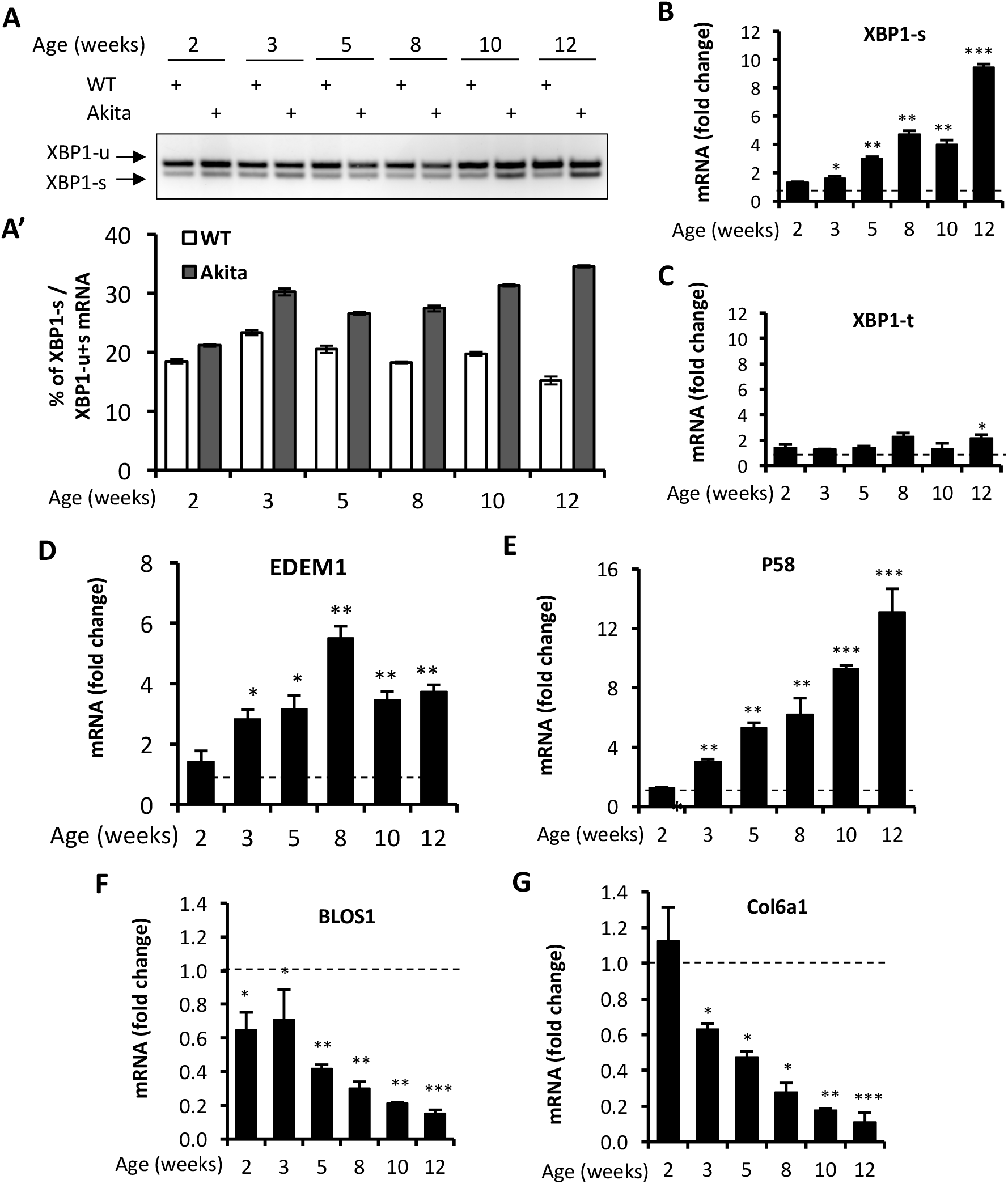
IRE1α RNase activity up-regulation in islets of Akita mice. **A-A’**. XBP1 mRNA levels were analyzed in islets isolated from Akita mice or age-matched C57B/6 mice at the indicated ages by RT-PCR, and the products were resolved by agarose gel electrophoresis. The full length (unspliced, XBP1-u) and spliced (XBP1-s) forms of XBP1 mRNA were indicated (A) and quantified (A’). Cyclophilin A mRNA (not shown) was used as an internal control. The data shown are representative of 3 independent experiments. **B-G**. mRNA levels for indicated genes were analyzed in islets isolated from Akita mice or age-matched C57B/6 mice by qRT-PCR. The results are expressed as the fold change over mRNA levels in respective age-matched controls (represented by the dashed line) and are representative of 3 independent experiments. * P < 0.05, ** P <0.01, and *** P <0.001. Bars indicate SEM.

### Treatment of IRE1α RNase inhibitor STF ameliorates diabetes in Akita mice

The results presented above in conjunction with previous observations that the overexpression of IRE-1α led to cell death in transfected cells (24, 46, 47) suggest that inhibiting IRE-1α may protect β-cells from Akita mutation-induced dysfunction and death and ameliorate the diabetic condition in Akita mice. We therefore investigated the effect of pharmacological inhibition of IRE-1α activity on β-cell and diabetic conditions in Akita mice. To test this, we treated Akita mice with a specific IRE-1α RNase inhibitor STF-083010 (STF, 15 mg/Kg of BW) (48) via intraperitoneal injection, starting at the age of 5-6 weeks for 6 weeks in which fasting blood glucose levels were at ∼280 mg/dL. While the saline-treated Akita mice further developed increasing hyperglycemia over time to reach up to 400mg/dL, the levels of blood glucose in the STF-treated Akita mice were gradually and significantly dampened over the 6-week period of treatment (Fig. 2A). Furthermore, the STF-treated Akita mice displayed a significantly improved glucose tolerance with lower peak glucose level and decreased AUC (area under the curve) following an intraperitoneal glucose bolus compared to their vehicle-treated counterpart (p<0.05; Fig. 2B, 2C). On the other hand, the STF- and vehicle-treated mice displayed no obvious differences in body weight (Fig. 2D) and insulin sensitivity (Fig. 2E-2F) suggesting that STF lowers blood glucose levels not through insulin sensitivity in the peripheral organs. Finally, in in vivo glucose stimulated insulin secretion assay, we observed a marked increase in serum insulin levels in STF-treated group compared to vehicle group; in particular, at 30 min after glucose injection (vehicle 0.2± 0.1 mg/ml vs. STF 1.5± 0.3 mg/ml; p<0.001) (Fig. 2G). Together, these results indicate that STF treatment significantly alleviates the diabetic conditions in Akita mice.

**Figure 2.**
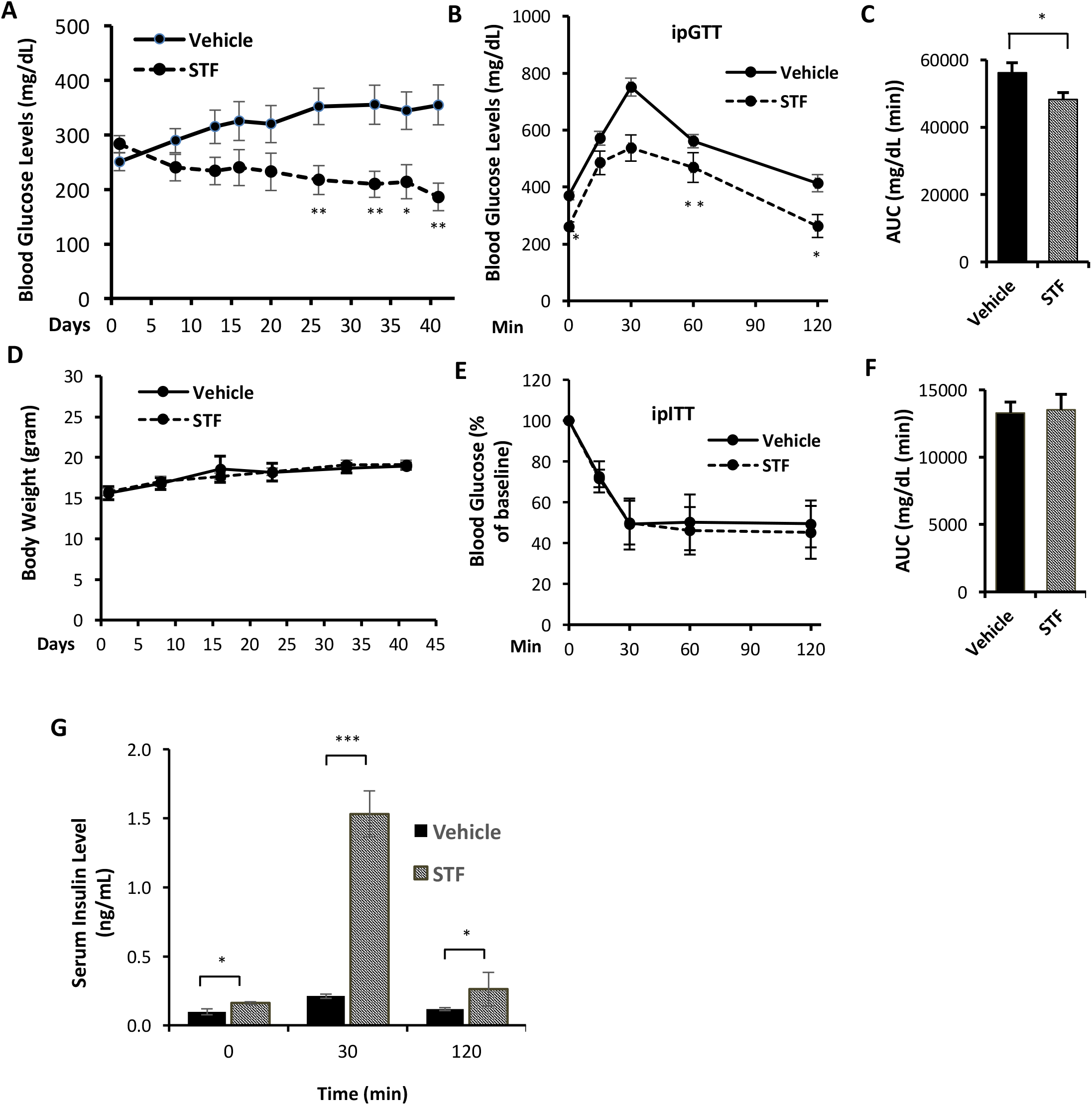
STF ameliorates diabetic conditions of Akita mice. **A**. Fasting blood glucose levels were measured in Akita mice treated with vehicle (n = 8) or STF (n = 7) at indicated time points. **B-C**. Glucose tolerance test. Blood glucose levels (B) measured at indicated time points after intraperitoneal injection of glucose (1.5g/kg body weight) following 6-h fasting and the AUC (area under the curve, C). **D**. Body weight of mice. **E-F**. Insulin tolerance test. Blood glucose levels (E) measured at indicated time points after intraperitoneal injection of insulin (0.75 IU/kg body weight) following 4-h fasting and the AUC (F). **G**. In vivo glucose-stimulated insulin secretion. Serum insulin levels measured at indicated time points after intraperitoneal injection of glucose (1.5g/kg body weight) following 6-h fasting. * P < 0.05, ** P <0.01, and *** P <0.001. Bars indicate SEM.

### STF treatment attenuates IRE1-α activity in islets of Akita mice

To investigate whether the STF amelioration of diabetic conditions in Akita mice is due to the inhibition of IRE1α activity in islets, we first examined the status of IRE1α-mediated *Xbp1* mRNA splicing in islets from STF-treated Akita mice. As shown in Fig. 3A, the level of *spliced XBP1* mRNA was significantly reduced in islets from Akita mice after 6-week treatment of STF compared to vehicle group. Corroborating these results, as assessed by qRT-PCR using Xbp1 splicing-specific primers, the increase in the spliced form of *Xbp1* mRNA levels was significantly attenuated by STF treatment, whereas the total *Xbp1* mRNA levels were only moderately affected by STF treatment (Fig. 3B, 3C). To further investigate whether STF treatment inhibits the IRE-1α activity, we evaluated the effect of STF treatment on the expression of target genes of XBP1. We observed that the mRNA levels of XBP1 target genes - *EDEM1* and *P58* –were strongly suppressed in the islets from STF-treated Akita mice relative to the vehicle-treated counterpart (Fig. 3D, 3E). Next, we evaluated the effect of STF on the RIDD target genes of IRE-1α. We found that the levels of the canonical RIDD target mRNAs Col61a and Blos1 were suppressed in the Akita islets; however, their levels were significantly reversed in islets of the STF-treated Akita mice (Fig. 3F, 3G). In addition, under ER stress, insulin 1 and insulin 2 mRNAs are known to be cleaved directly by IRE1-α RNase activity and are therefore RIDD target mRNAs in β-cells (24, 49). Consistent with this, both insulin 1 and insulin 2 mRNAs were down-regulated in Akita islets (Fig. 3H, 3I). However, STF treatment significantly reversed their expression in the Akita islets (Fig. 3H, 3I). Interestingly, we notice that although STF is known to inhibit IRE-1α activity only, our results revealed that STF also suppressed the Akita mutation-induced increase in the mRNA levels of *ATF4* and *CHOP*, key components in PERK pathway (Fig. S2A, S2B), and of *Bip* (Fig. S2C), which is controlled by XBP1/ATF6 heterodimer and hence also a target of ATF6 pathway. These effects of STF on PERK and ATF6 pathways could be due to cross-talks among the branches of UPR under in vivo conditions (50, 51). Together, our results reveal that STF treatment suppresses Akita mutation-induced IRE-1α activation in islets of Akita mice. Notably, the correction of glucose homeostasis defects in Akita mice by STF is associated with the reversal of hyperactivated IRE1α activity to normal level and not the abolishment of basal IRE-1α activity (Fig. 3A-B).

**Figure 3.**
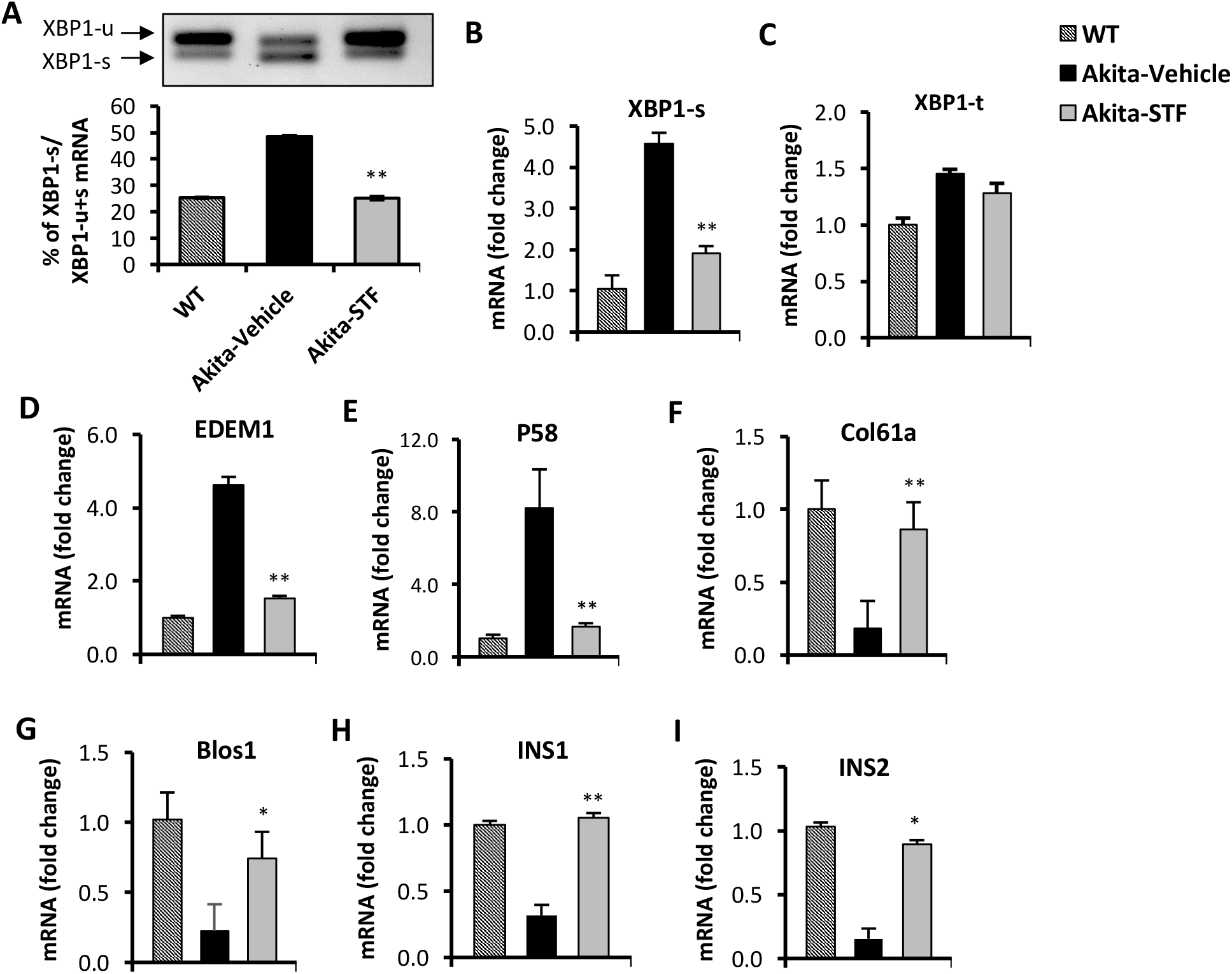
STF inhibits IRE1α RNase activity. **A**. XBP1 mRNA levels were analyzed in islets isolated from Akita mice treated with STF or vehicle by RT-PCR and the products were resolved by agarose gel electrophoresis. The full length (unspliced, XBP1-u) and spliced (XBP1-s) forms of XBP1 mRNA were indicated and quantified by ImageJ program. Cyclophilin A mRNA (not shown). **B-I**. mRNA levels for indicated genes were analyzed in islets isolated from Akita mice treated with STF or vehicle by qRT-PCR. The results are expressed as fold change and are representative of 3 independent experiments. * P < 0.05, ** P <0.01, and *** P <0.001 compared to Akita-vehicle group. Bars indicate SEM.

### STF promotes β-cell viability in Akita mice

As the development of diabetes in Akita mice is associated with gradual β-cell loss and IRE-1α is activated before the onset of diabetes in Akita mice, we investigated whether the STF improvement of diabetic conditions is associated with the protection of islet β-cells in the Akita mice. To this end, we examined the status of β-cell mass and survival in STF-treated Akita mice. In pancreatic sections, Akita mice not only exhibited reduced β-cell mass but also significantly decreased insulin staining intensity in existing β-cells (Fig. 4A, 4B, 4D, 4E), indicative of β-cell loss and dysfunction. In contrast, the STF-treated Akita mice had approximately twice the β-cell mass and significantly higher insulin staining intensity compared to that in vehicle-treated mice (Fig. 4B-4E). On the other hand, the α cell numbers per islet between STF-treated and vehicle-treated Akita groups remain comparable, as marked by glucagon immunostaining (Fig. 4A-4C). Consistent with these results, total pancreatic insulin content as quantified by ELISA was markedly higher in STF-treated Akita mice than in their vehicle-treated counterparts (Fig. 4F). To determine whether the increase in β-cell mass by STF could be attributed to an inhibition of islet cell apoptosis, we assessed apoptosis in the Akita mouse pancreatic islets by performing TUNEL (terminal deoxynucleotidyl transferase dUTP nick end labeling) staining, a marker for apoptosis. An increase in TUNEL ^+^ insulin^+^ cells was observed in the vehicle-treated Akita mice relative to wild-type mice (Fig. 4G-J). However, treatment with STF considerably reduced TUNEL staining in the islet β-cells of Akita mice to a level comparable to that of wild-type islets (Fig. 4G-J). Corroborating these findings, we observed that treatment with STF significantly reduced the number of CASP^+^ (a critical protein in the execution of apoptosis) insulin^+^ cells of Akita mice compared to the vehicle-treated counterpart (Fig. S3A-D). In contrast, STF treatment appeared to exhibit no effect on β-cell proliferation in that the percentage of Ki67^+^ insulin^+^ cells appeared to be similarly rare in both vehicle and STF groups (Fig. S4). Lastly, expression of the apoptotic effector genes BAX and Bak1, as well as expression of the pro-apoptotic inducer gene p53 and a negative cell-cycle regulator p21, were significantly suppressed in the islets of STF-treated Akita mice (Fig. S5A-D). In sum, these results indicate that inhibition of increased IRE1-α activity by STF suppresses β-cell apoptosis in Akita mice which in turn leads to a preservation of β cell mass.

**Figure 4.**
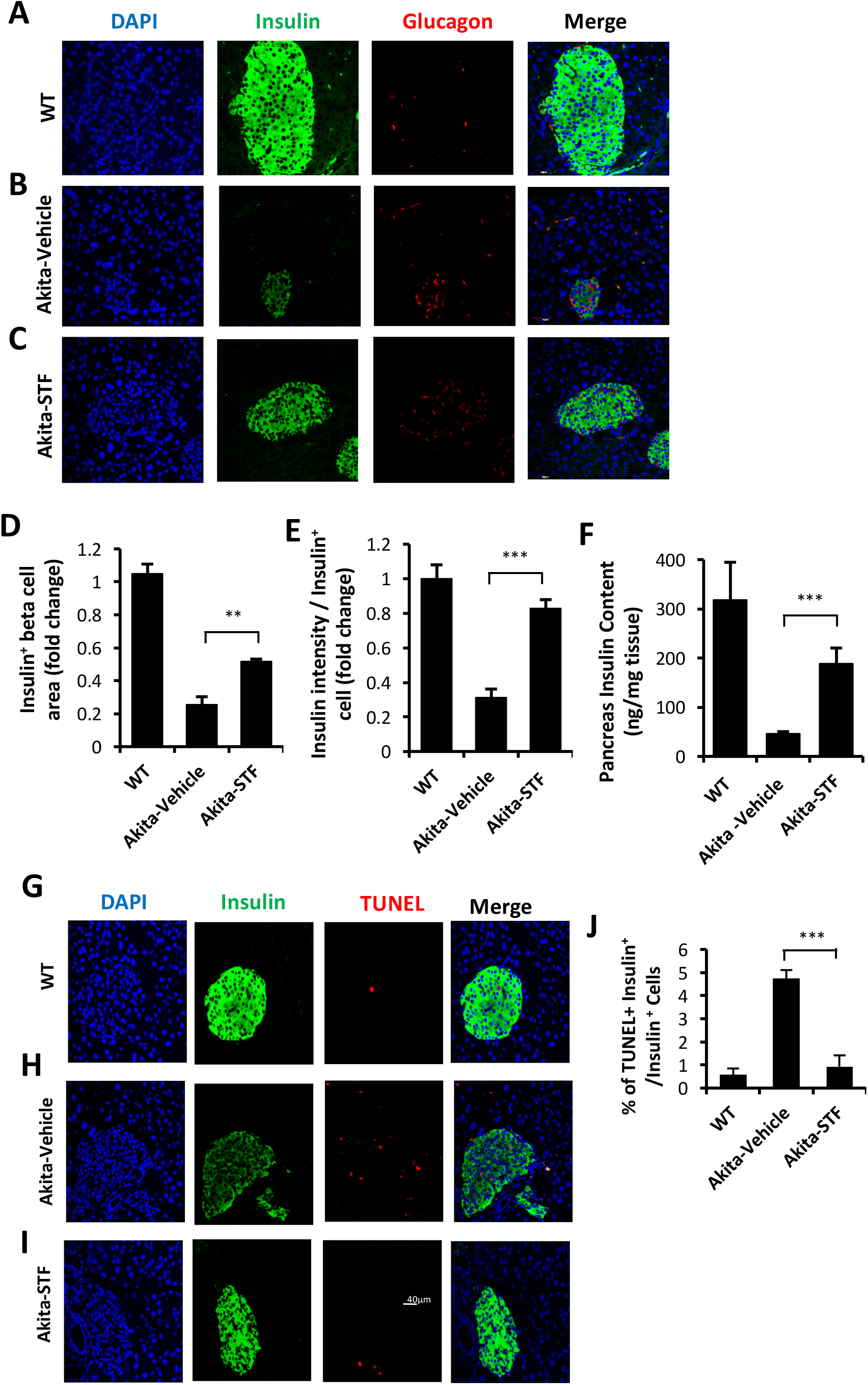
STF preserves β-cell mass and viability and suppresses β-cell apoptosis in Akita mice. **A-C**. Immunofluorescence staining of pancreatic sections. Pancreases were sectioned and slides were stained with anti-insulin antibody (green, β-cell marker), anti-glucagon antibody (red, α-cell marker), and DAPI (blue). Slides were imaged with an Olympus FV1000 confocal microscope. **D**. Quantification of insulin^+^ β-cell area. Total area of all islets per section was calculated from a total of six sections for each of three mice using insulin^+^ cells to demarcate islet β-cells and normalized with that for C57B/6 mice designated as 1. **E**. Insulin staining intensity. The average insulin staining intensity was quantified using ImageJ and normalized with that for C57B/6 mice designated as 1. **F**. Insulin content measurement by ELISA as detailed in Methods and Materials. **G-I**. TUNEL staining in pancreatic sections. Pancreatic sections were stained with anti-insulin antibody (green, β-cell marker), TUNEL (red, cell death), and DAPI (blue). Slides were imaged with an Olympus FV1000 confocal microscope. **J**. Quantification of percentage of TUNEL^+^ insulin^+^ β-cells/insulin^+^ cells. At least 50 islets were counted for each group. Data are the mean± SEM. * P < 0.05, ** P <0.01, and *** P <0.001.

### STF improves β-cell function in Akita mice

We interrogated whether STF also improves β-cell function. Indeed, our observations that STF heightened the production of insulin in β-cells (Fig. 4C-4F) and STF treatment increased serum insulin levels (Fig. 2G) suggest an improvement in β-cell function in mice with STF treatment. To further investigate the protective effect of STF on β-cell function, we first assessed the glucose-stimulated insulin secretion in islets isolated from vehicle or STF-treated mice. As shown in Fig. 5A, insulin secretion was significantly higher under both basal (2.5 mM glucose concentration) and stimulated (16.7 mM glucose concentration) conditions in islets isolated from the STF-treated Akita mice compared to vehicle-treated counterpart. Second, it was reported that high glucose and the ensuing ER stress in β-cells down-regulate the expression of β-cell-specific transcription factors including PDX1 and MafA (27, 52-55). Transcription factors such as PDX1, MafA, Nkx6.1, and NeurD1 are not only required for proper β-cell development but are also essential for the maintenance of normal β-cell function including insulin mRNA transcription (56-58). We therefore investigated the effect of Akita mutation on the expression levels of these transcription factors in β-cells and the impact of STF treatment on their expression levels in Akita β-cells. As shown in Figures 5B-5E, the mRNA levels of *Pdx1, MafA, NeurD1*, and *Nkx6.1* were significantly down-regulated in islets of Akita mice compared to age- and sex-matched WT mice. STF treatment markedly reversed the reduction in the expression of these genes in islets from treated Akita mice.

**Figure 5.**
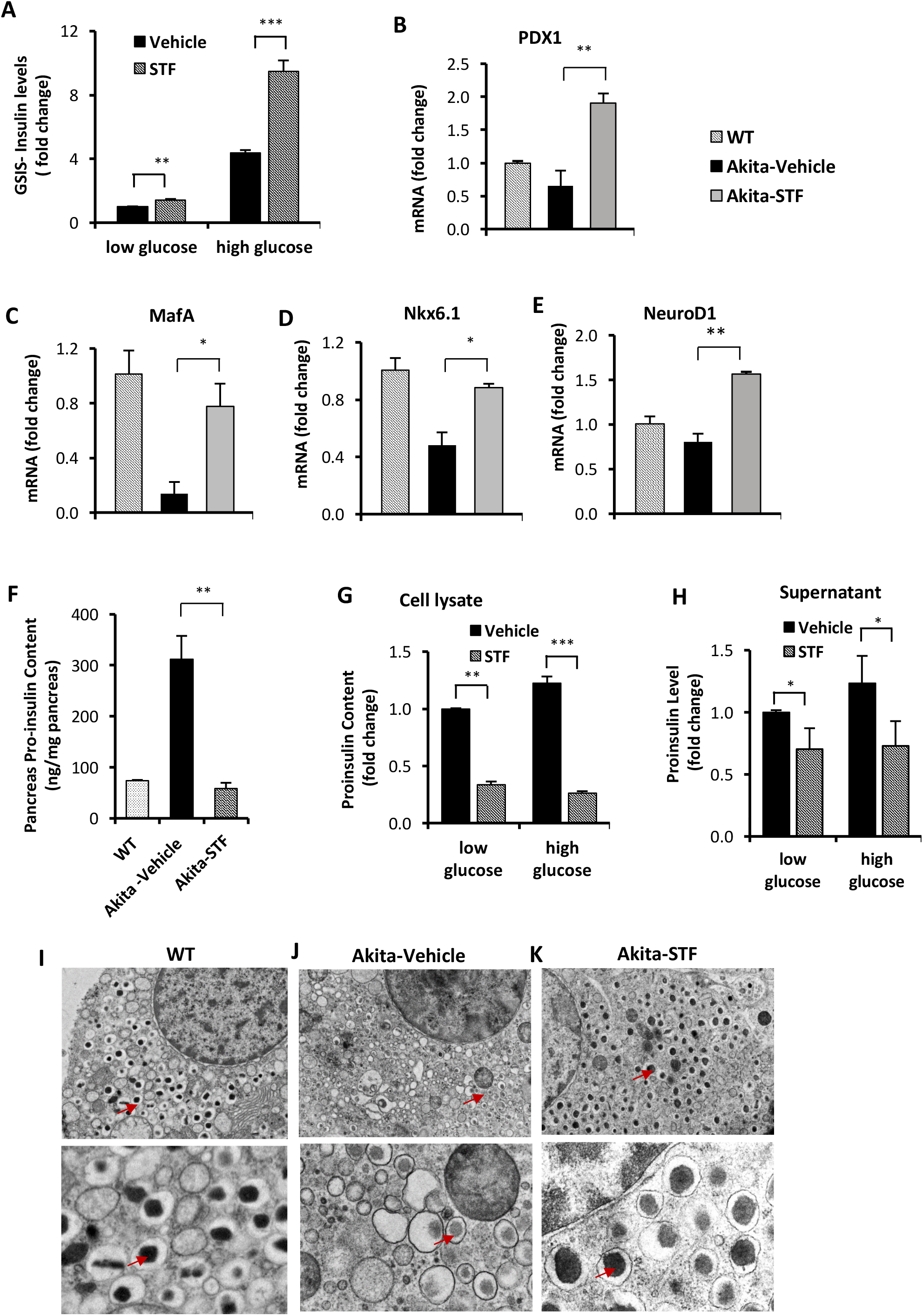
STF improves Akita islet β-cell function. **A**. Glucose-stimulated insulin secretion of 20 islets isolated from Akita mice treated with STF or vehicle and incubated with 2.5 mM and 16.7 mM glucose. Secreted insulin was measured by ELISA. The data was presented as fold change and normalized with total protein concentration, with the amount of insulin secreted in response to 2.5 mM glucose from vehicle-treated group set to 1.0. **B-E**. mRNA levels for indicated genes were analyzed in islets isolated from Akita mice treated with STF or vehicle by qRT-PCR. The results are expressed as fold change and are representative of 3 independent experiments. **F**. Proinsulin content measurement by ELISA as detailed in Methods and Materials. **G-H.** Proinsulin content and secretion measurement. 20 islets isolated from Akita mice treated with STF or vehicle were incubated with 2.5 mM and 16.7 mM glucose. Secreted proinsulin was measured by ELISA. Proinsulin content measurement by ELISA as detailed in Methods and Materials. The data was presented as fold change and normalized with total protein concentration, with the amount of proinsulin in response to 2.5 mM glucose from vehicle-treated group set to 1.0. **I-K**. Ultrastructure of β-cells in islets isolated from Akita treated with STF by transmission electron microscopy. Images at the top panel and the bottom panel were taken at 3,000x and 10,000x, respectively. Arrows point to dark mature insulin granules.

Akita mutant proinsulin protein tends to not only misfold on its own but also forms heterogeneous complex with wild-type (WT) proinsulin molecules, thus entrapping both Akita and WT proinsulin in the ER (59-61) and limiting bioactive insulin production and secretion, leading to ER stress and β-cell dysfunction and death (39, 62). The disturbed ER environment in turn further exacerbates the proinsulin misfolding problem (9). We therefore investigated whether the increased insulin production (Figs. 2G, 4C-F, 5A) seen in STF treatment is associated with reduced proinsulin levels. First, we analyzed the effect of STF treatment on the proinsulin content levels in pancreas from STF- or vehicle-treated Akita mice. As expected, proinsulin content was dramatically and significantly increased in the Akita mouse pancreas relative to WT pancreas, but was completely suppressed to normal level in the pancreas of the STF-treated Akita mice (Fig. 5F). Of note, the STF effect on proinsulin is opposite to that seen for insulin whereby the insulin content was strongly reduced in the pancreatic islets of Akita mice relative to control mice and STF treatment significantly raised the blunted insulin level to close to normal level (Fig. 4F). While we interpret the STF suppression of proinsulin level in Akita islets as the outcome of improved ER environment by STF which permits more proinsulin conversion to mature insulin; however, it is also possible that this effect is mediated through increased proinsulin release from β cells. To address this possibility, we analyzed proinsulin levels in islets that have been isolated from vehicle- or STF-treated Akita mice under either basal (2.5 mM) or high (16.7 mM) glucose concentration. As shown in Figures 5G and5H, both proinsulin content (cell lysate) and secretion (supernatant) were attenuated in the islets isolated from the STF-treated Akita mice compared to vehicle-treated mice, under both basal (2.5 mM glucose) and stimulated (16.7 mM glucose) conditions, thereby indicating that STF suppresses total proinsulin content without affecting proinsulin secretion from the pancreatic islets.

Finally, as mature insulin is formed from proinsulin processed in the Golgi complex and stored in secretory granules for release, we examined the effect of STF on the insulin secretory granules and the ultrastructure of islet β-cells using transmission electron microscopy. Our results showed that whereas there was markedly reduced number of dark dense-core insulin granules (mature insulin granules), increase in number of light or “gray” dense-core granulels (inmature insulin granules) in Akita β-cells compared to WT β-cells (Fig. 5I-J), β-cells in the STF-treated Akita mice exhibited a marked increase in the number of dark electron-dense core insulin granules and (Fig. 5K), similar to those seen in the WT islets (Fig. 5I). Together, these results demonstrate that STF normalization of IRE-1α activity facilitates insulin granule formation.

### STF suppresses ER stress-related inflammation and oxidative stress in the islets of Akita mice

ER stress has been shown to cause and potentiate inflammation and oxidative stress that cooperatively contribute to ER stress-mediated cell death (63). We therefore investigated whether STF treatment affects these processes in the Akita islets. We first analyzed the expression levels of ER stress-associated proinflammatory cytokines and found that mRNA levels of the cytokine genes IL-1β, IL6, and TNF were increased in the Akita pancreatic islets compared to WT islets (Figs. 6A-6C). In agreement with an occurrence of ER stress-associated low-grade inflammation, we also detected increased transcript levels of MCP1 and CD68 (Figs. 6D, 6E), markers that are highly expressed in tissue monocytes and macrophages, respectively. Strikingly, STF corrected the mRNA levels of these genes to normal levels in the pancreatic islets of Akita mice (Figs. 6A-6E).

**Figure 6.**
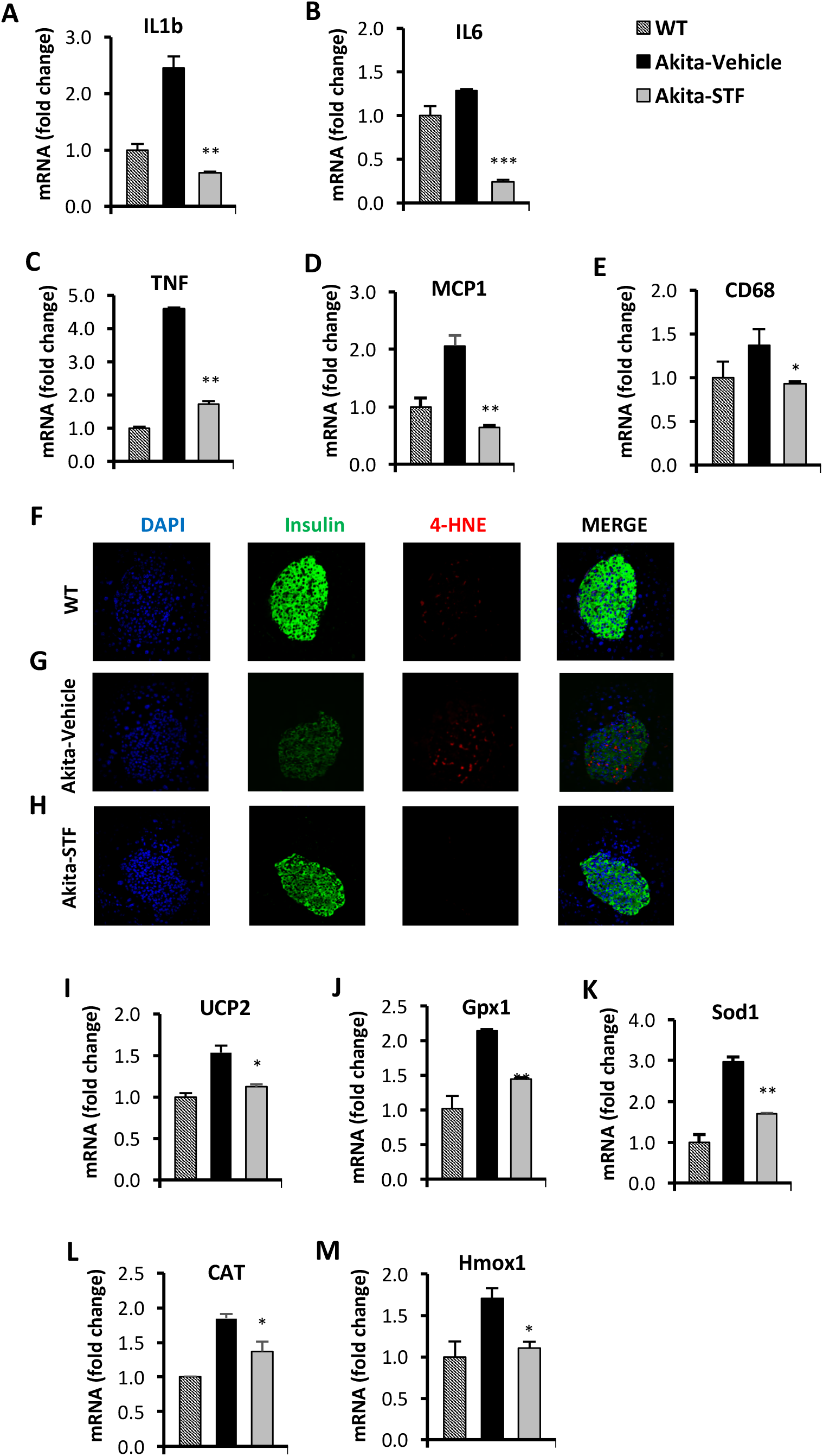
STF attenuates the ER stress-associated inflammation and oxidative stress. **A-E.** mRNA levels for indicated genes involving inflammation were analyzed in islets isolated from Akita mice treated with STF or vehicle by qRT-PCR. The results are expressed as the fold-increase over mRNA levels and are representative of 3 independent experiments. **F-J**. mRNA levels for indicated anti-oxidant genes were analyzed in islets isolated from Akita mice treated with STF or vehicle by qRT-PCR. The results are expressed as fold change and are representative of 3 independent experiments. * P < 0.05, ** P <0.01, and *** P <0.001 compared to Akita-vehicle group. Bars indicate SEM.

Next, we assessed the effect of STF on oxidative stress in Akita islets. In Akita mice, we found an obvious nuclear accumulation of the lipid peroxidation product 4-hydroxynonenal (4-HNE), a marker of oxidative stress, in the pancreatic islets of the vehicle-treated Akita mice compared to WT mice (Fig. 6F, 6G). Similarly, we observed an up-regulation of the mRNA levels of several antioxidant genes, including those encoding the mitochondrial uncoupling protein 2 (UCP2), glutathione peroxidase 1 (Gpx1), superoxide dismutase 1 (Sod1), catalase (CAT), and heme oxygenase 1 (Hmox1) in the pancreatic islet cells of Akita mice compared to WT mice (Fig. 6I-6M), reflecting a compensatory mechanism of anti-oxidation through the antioxidant gene up-regulation (64, 65). Notably, the nuclear accumulation of 4-HNE was abolished in the pancreatic islets of Akita mice treated with STF (Fig. 6G, 6H). Likewise, STF treatment significantly ameliorated the up-regulated expression of the antioxidant genes (Fig. 6I-6M).

### Ire1-α RNase inhibitor 4μ8C ameliorates diabetic conditions in Akita mice

To ascertain that STF improves diabetic conditions of Akita mice via the inhibition of IRE1α, we used another structurally distinct IRE1α RNase inhibitor 4μ8C (66) for the efficacy study. Intraperitoneal injection of 4μ8C (10 mg/kg of body weight, once daily) for 5 weeks improved the fasting blood glucose levels in Akita mice while vehicle-treated Akita mice showed a progressive rise in blood glucose level. At the end of the 5-week treatment, blood glucose levels in the 4μ8C-treated Akita mice were nearly normalized (Fig. 7A). 4μ8C treatment also significantly improved glucose tolerance in Akita mice (Fig. 7B-7C), with no significant difference in body weight (Fig. S6) and insulin tolerance (Fig. 7D), compared to vehicle-treated Akita mice. The normalization of blood glucose levels by 4μ8C treatment correlated with a marked increase in serum insulin levels and pancreatic insulin content in Akita group treated with 4μ8C (Figs. 7E, S7). Moreover, 4μ8C treatment significantly preserved the β-cell area and restored insulin staining intensity in β-cells (Fig. 7F-7J), thereby indicating the 4μ8C correction of the β-cell loss and dysfunction in Akita mice. Furthermore, as expected, we observed that 4μ8C treatment significantly suppressed the Akita mutation-induced increase in the ratio of spliced *XBP1* mRNA (Fig. S8A-B). Consistent with the alleviation of *XBP1* splicing, 4μ8C attenuated the heightened mRNA levels of XBP1-s target genes Grp94, Bip and P58 in Akita islets (Fig. S8C-E) and significantly reversed the repressed levels of RIDD target mRNAs Blos1, Col6a1, and INS1 in Akita mice to near WT levels (Fig. S8F-H). Lastly, 4μ8C attenuated the increased levels of PERK pathway genes ATF4 and CHOP in Akita mouse pancreatic islets (Fig. S8I-J). Together, we conclude that inhibition of Ire1α RNase activity is key in correcting the diabetic conditions of Akita mice.

**Figure 7.**
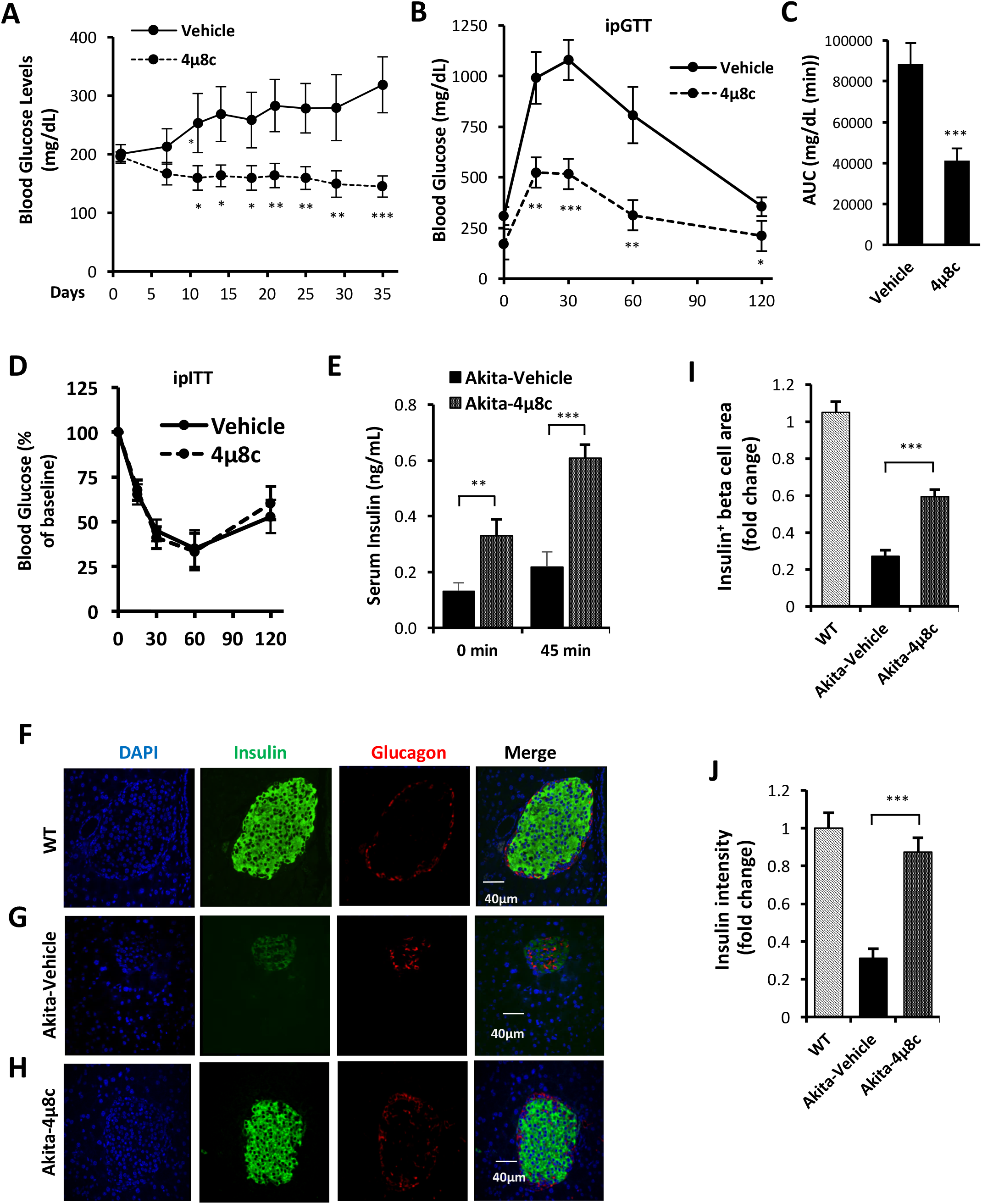
4μ8C ameliorates diabetic conditions of Akita mice and inhibits IRE1α RNase activity. **A**. Fasting blood glucose levels were measured in Akita mice treated with vehicle (n = 7) or 4μ8C (n = 6) at indicated time points. **B-C**. Glucose tolerance test. Blood glucose levels (B) measured at indicated time points after intraperitoneal injection of glucose (1.5g/kg body weight) following 6-h fasting and the AUC (area under the curve, C). **D**. Insulin tolerance test. Blood glucose levels measured at indicated time points after intraperitoneal injection of insulin (0.75 IU/kg body weight) following 4-h fasting. **E**. In vivo glucose-stimulated insulin secretion. Serum insulin levels measured at indicated time points after intraperitoneal injection of glucose (1.5g/kg body weight) following 6-h fasting. **F-H**. Immunofluorescence staining of pancreatic sections. Pancreases were sectioned and slides were stained with anti-insulin antibody (green, β-cell marker), anti-glucagon antibody (red, α-cell marker), and DAPI (blue). Slides were imaged with an Olympus FV1000 confocal microscope. **I**. Quantification of insulin^+^ β-cell area. Total area of all islets per section was calculated from a total of six sections for each of three mice using insulin^+^ cells to demarcate islet β-cells and normalized with that for C57B/6 mice designated as 1. **J**. Insulin staining intensity. The average insulin staining intensity was quantified using ImageJ and normalized with that for C57B/6 mice designated as 1. * P < 0.05, ** P <0.01, and *** P <0.001. Bars indicate SEM.

## DISCUSSION

In this study, we observed that IRE-1α activity was progressively up-regulated in the islets of Akita mice in an age-dependent fashion and that the increased IRE1α activity predates the onset of diabetes in Akita mice. Importantly, we showed that the inhibition of IRE-1α RNase activity markedly ameliorated the diabetic conditions and protected β-cell viability and function in Akita mice, thus revealing IRE-1α as an important target in β-cell protection and diabetes therapy. Using two pharmacological IRE-1α RNase inhibitors STF and 4μ8c, we showed that fasting blood glucose level was reduced while glucose tolerance and serum insulin were improved in the Akita mice with STF or 4μ8c administration. Concomitantly, the decline in pancreatic β-cell number and function in Akita mice was restored in mice treated with STF or 4μ8c. Taken together, our data indicate that the inhibition of Ire1α RNase activity plays a central role in ameliorating β-cell dysfunction and death, which culminates in the alleviation of hyperglycemia in Akita mice,

In further pursuit of how the inhibition of Ire1α RNase activity protects against β-cell dysfunction and loss in Akita mice, we examined the effect of STF on ER stress/UPR. As expected, the level of Ire1α RNase activity was reduced in in the STF-treated Akita pancreatic islet cells. Interestingly, we observed that Ire1α inhibition also suppressed insulin misfolding-induced activation of PERK and ATF6 pathways of UPR, which likely reflects the cross talk among different UPR pathways (67, 68). ER stress crosstalks with and exacerbates inflammation and oxidative stress each other (63). As a corollary, the levels of inflammation and oxidative stress were up-regulated in Akita islet cells but suppressed by the treatment of Ire1α inhibitor. In all, our data reveal the amelioration of ER stress and downstream inflammation and oxidative stress as the underlying mechanisms of the protection of β-cell dysfunction and demise in Akita mice by inhibiting Ire1α activity.

Previous studies have reported that imatinib and similar tyrosine kinase inhibitors exhibit β cell protection by inhibiting IRE1α kinase activity, either directly or through an intermediary factor, leading to the attenuation of IRE1α RNase activity (25, 69, 70). However, given the promiscuous nature of kinase inhibitors which generally target multiple kinases, it is possible that IRE1α might not be the sole kinase target of imatinib and related tyrosine kinase inhibitors. Furthermore, in addition to acting as kinase inhibitor, imatinib has also been reported to serve as a partial agonist of peroxisome proliferator-activated receptor gamma (PPARγ) (71, 72) as well as a modulator of autophagy (73-75), both of which exhibit β cell protection. PPARγ was shown to protect β cells in diabetes by suppressing ER stress (76, 77). The role of autophagy on β cell function and viability has also been well documented (78, 79). Therefore, the effect of imatinib and related tyrosine kinase inhibitors on β cell protection is likely the outcome of acting on multiple factors including IRE1α inhibition. However, how significantly IRE1α inhibition contributes to the improvement of diabetes or whether IRE1α inhibition alone, in particular, the sole inhibition of IRE1α RNase function as IRE1α kinase domain also plays key role in many biological functions (80), is sufficient to ameliorate diabetic condition is unclear. Our current work, on the other hand, provides clear evidence that inhibiting IRE1α RNase activity alone with two different IRE1α RNase inhibitors STF and 4μ8c is sufficient to significantly improve the diabetic condition and β cell function and health in Akita mice.

IRE1α has been documented to serve an important modulatory role in multiple physiological contexts including β cell function, growth and survival, which poses a question on whether IRE1α inhibition would improve or even exacerbates β cell and diabetic conditions in an ER stress-related situation such as proinsulin misfolding Akita diabetes model. As our results showed, in contrast to IRE1α β cell deletion which leads to β cell functional impairment and growth defects, pharmacological inhibition of IRE1α markedly ameliorated diabetic condition and improved β cell mass and function in Akita mice. We note that pharmacological inhibition of IRE1α did not totally abolish the IRE1α function as genetic knockout would do; instead, IRE1α inhibitors simply reversed hyperactivated IRE1α back to normal or basal level of activity as shown by our data. The normalization of IRE1α activity by inhibitors in turn reversed IREα hyperactivation-induced diabetic conditions in Akita mice, without the unwanted side effects associated with IREα knockout. Our findings therefore highlight the notion that the normalization (not elimination) of IRE1α activity as the key to the success of the therapeutic use of pharmacological inhibitors on proteins with physiologically important but pathologically heightened activity.

In summary, our studies showed that pharmacological IRE1α inhibition preserves β-cells and prevents the development of diabetes in insulin protein misfolding-causing Akita mice. This protection is associated with significant increase in the number of β-cells through the attenuation of apoptosis and the preservation of β-cell function including basal and glucose-stimulated insulin secretion. IRE1α inhibitors achieved these effects through the suppression of ER stress-induced excessive activation of UPR. These findings may offer an effective therapeutic strategy for MIDY patients. In addition, as ER stress and insulin misfolding are well established in their roles in β cell dysfunction and demise in type 2 diabetes, IRE1α inhibition may well be considered for the treatment of type 2 disease.

## ACKNOWLEDGMENTS

This work was supported by Oklahoma Center for the Advancement of Science and Technology and National Institutes of Health (Grants GM103636, DK108887, DK116017) to W.W. Research reported in this publication was supported in part by the National Cancer Institute Cancer Center Support Grant P30CA225520 and the Oklahoma Tobacco Settlement Endowment Trust contract awarded to the University of Oklahoma Stephenson Cancer Center and used the Biospecimen and Tissue Pathology and Molecular Biology and Cytometry Research of the CCSG Shared Resources, Imaging Core of NIH COBRE (5P30GM103636), and Histology and Imaging Cores of Diabetes COBRE (5P30GM122744).

## Author Contributions

O.H.-P., V.E., R.B.U., D.W., I.R. generated research data. H.-Y.L. designed the research project, contributed to discussion, and reviewed/edited the manuscript. W.W. conceived, initiated, and designed the research project, reviewed the data, and wrote the manuscript. W.W. is the guarantor of this work and, as such, had full access to all the data in the study and take responsibility for the integrity of the data and the accuracy of the data analysis.

---

**Supplemental Figure 1.**
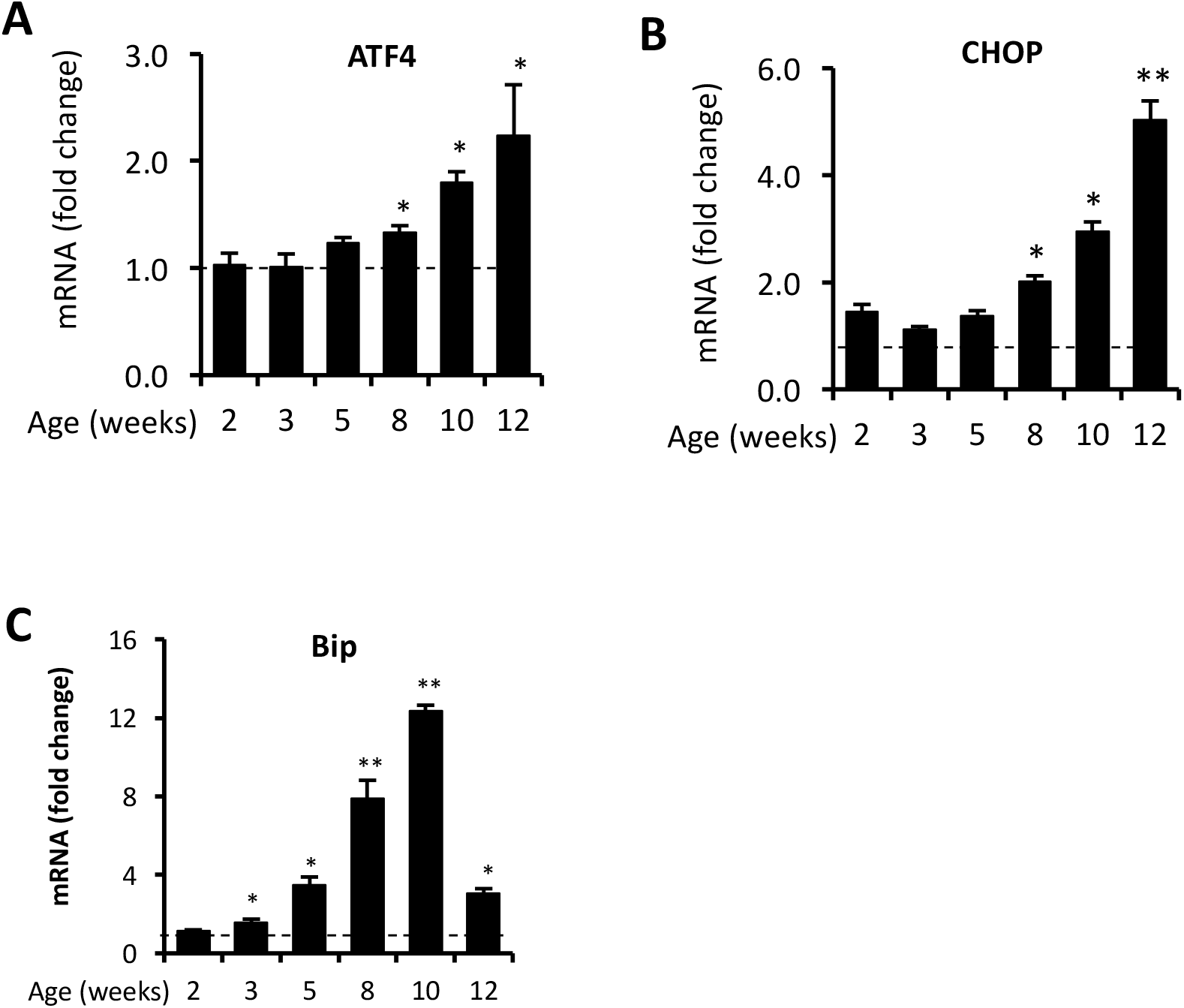
**A-C**. mRNA levels for indicated genes were analyzed in islets isolated from Akita mice or age-matched C57B/6 mice by qRT-PCR. The results are expressed as the fold change over mRNA levels in respective age-matched controls (represented by the dashed line) and are representative of 3 independent experiments. * P < 0.05, ** P <0.01, and *** P <0.001. Bars indicate SEM.

**Supplemental Figure 2.**
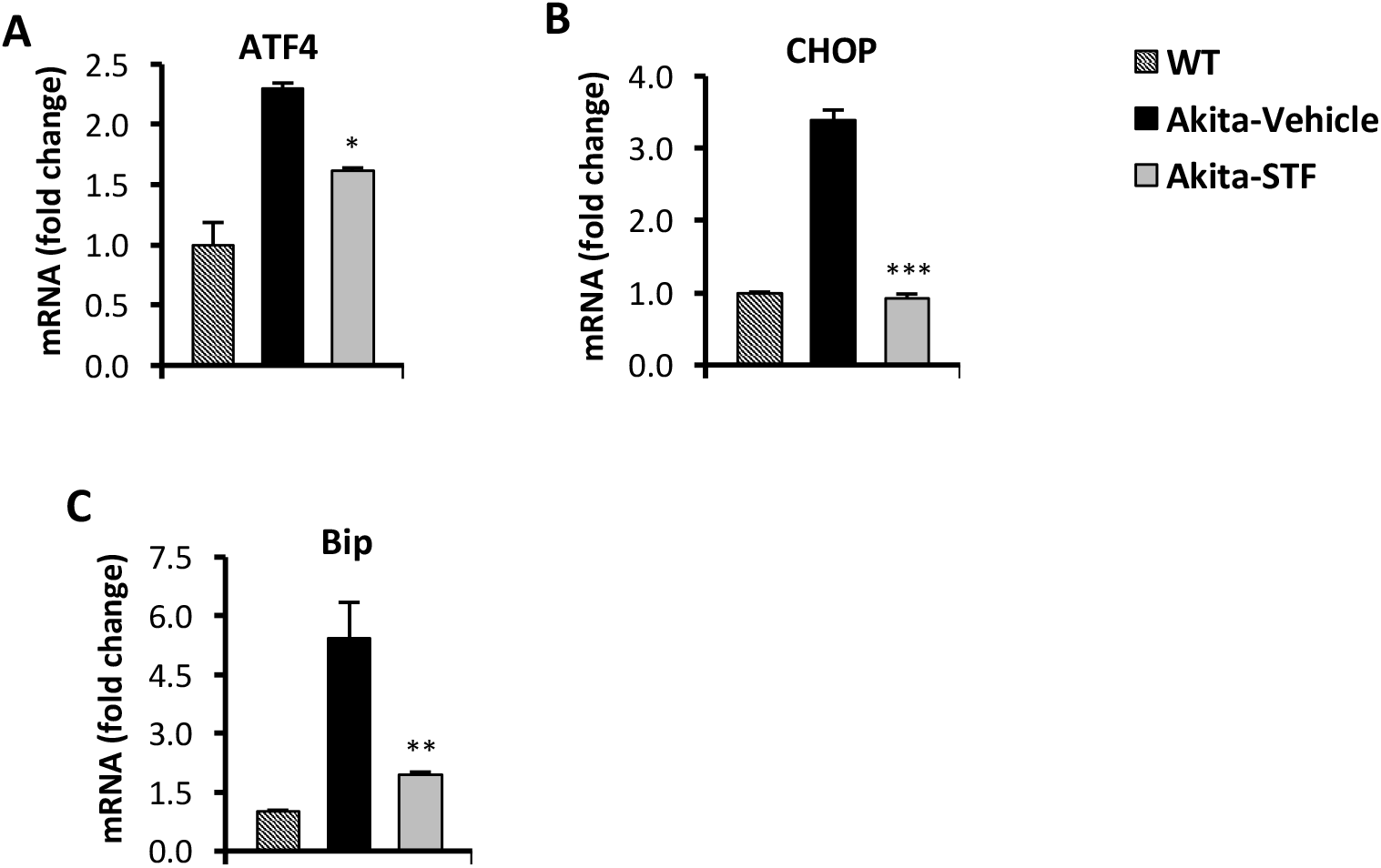
**A-C**. mRNA levels for indicated anti-oxidant genes were analyzed in islets isolated from Akita mice treated with STF or vehicle by qRT-PCR. The results are expressed as fold change and are representative of 3 independent experiments. * P < 0.05, ** P <0.01, and *** P <0.001 compared to Akita-vehicle group. Bars indicate SEM.

**Supplemental Figure 3.**
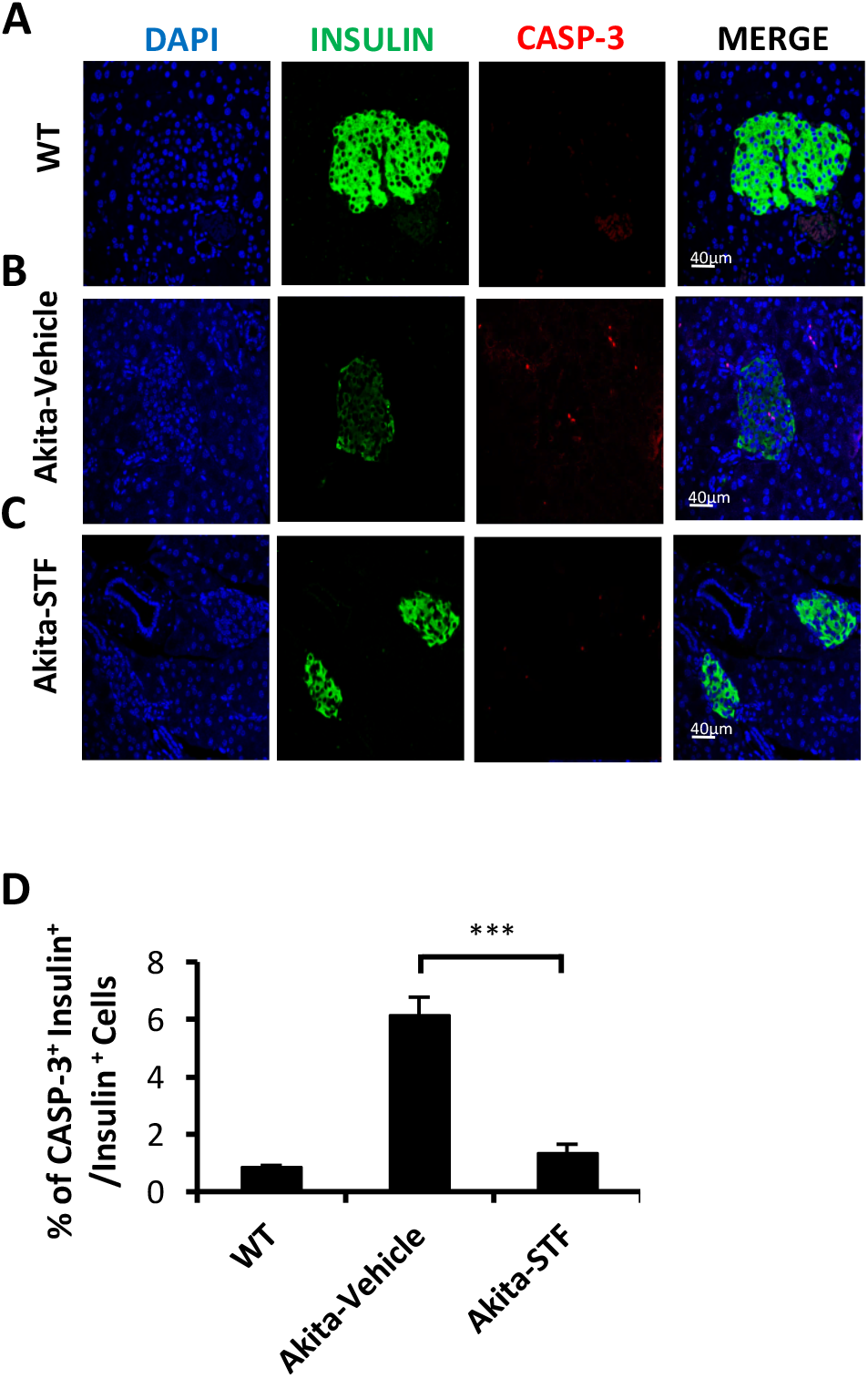
**A-C**. Immunofluorescence staining of pancreatic sections. Pancreases sections were stained with an-Casp3 antibody (red), anti-insulin antibody (green, β-cell marker), and DAPI (blue). Slides were imaged with an Olympus FV1000 confocal microscope. **D**. Quantification of percentage of Casp3^+^ insulin^+^ β-cells/insulin^+^ cells. At least 50 islets were counted for each group. Data are the mean±SEM. *** P <0.001.

**Supplemental Figure 4.**
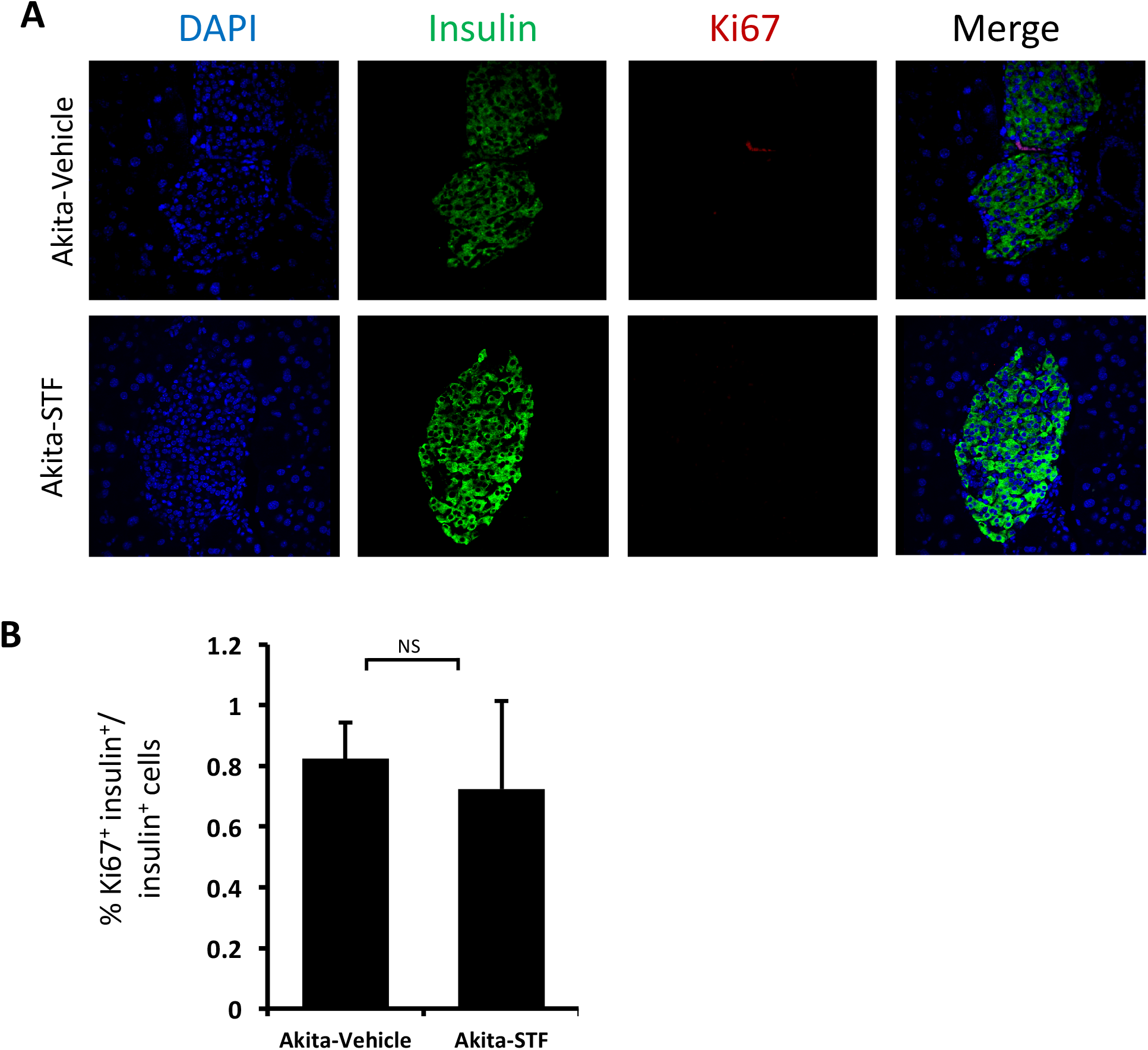
**A.** Immunofluorescence staining of pancreatic sections. Pancreases sections were stained with an-Ki67 antibody (red), anti-insulin antibody (green, β-cell marker), and DAPI (blue). Slides were imaged with an Olympus FV1000 confocal microscope. **B**. Quantification of percentage of Ki67^+^ insulin^+^ β-cells/insulin^+^ cells. At least 50 islets were counted for each group. Data are the mean±SEM. NS, P >0.05.

**Supplemental Figure 5.**
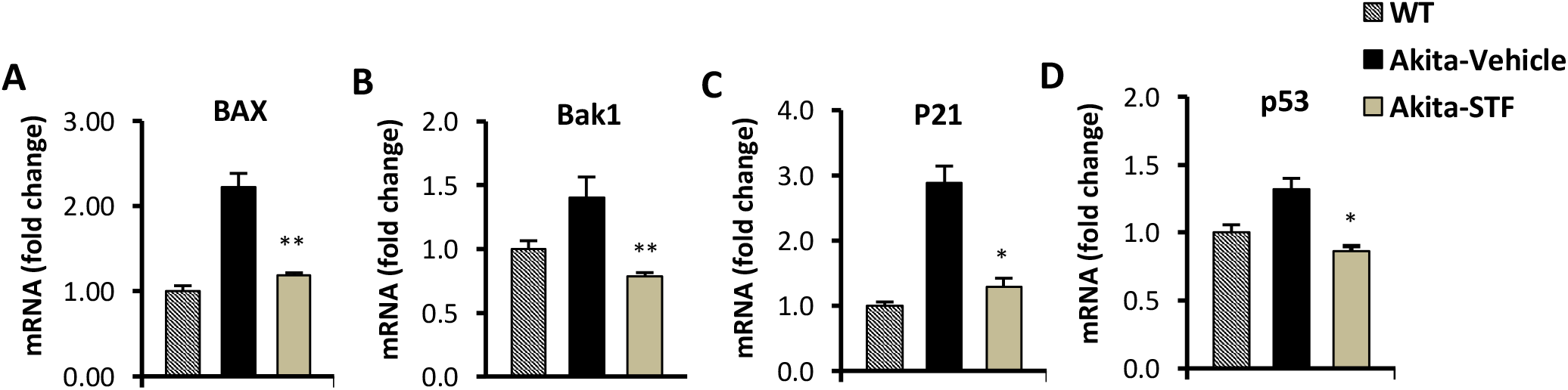
**A-D**. mRNA levels for indicated anti-oxidant genes were analyzed in islets isolated from Akita mice treated with STF or vehicle by qRT-PCR. The results are expressed as fold change and are representative of 3 independent experiments. * P < 0.05, ** P <0.01, and *** P <0.001 compared to Akita-vehicle group. Bars indicate SEM.

**Supplemental Figure 6.**
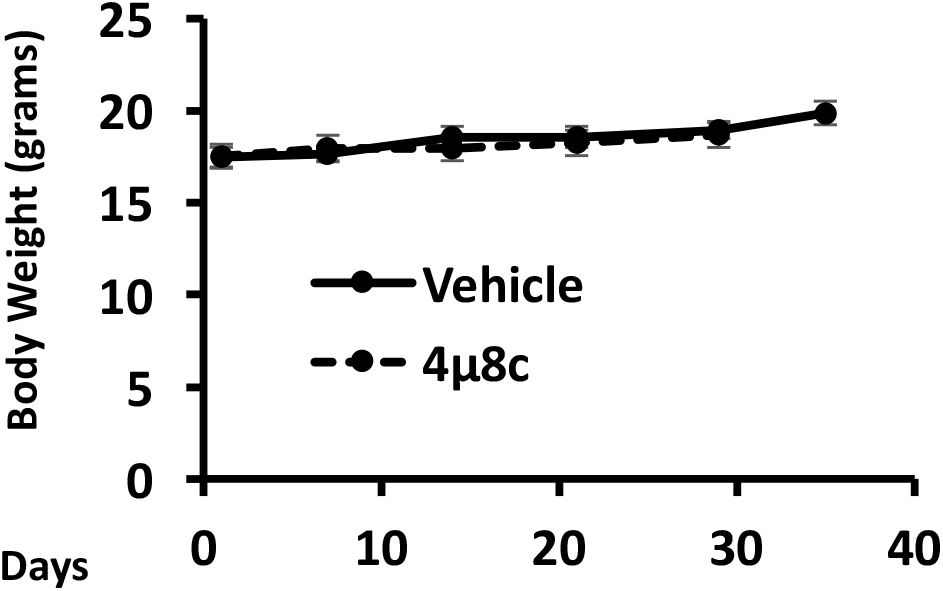
Body weight of Akita mice treated with vehicle or 4μ8C.

**Supplemental Figure 7.**
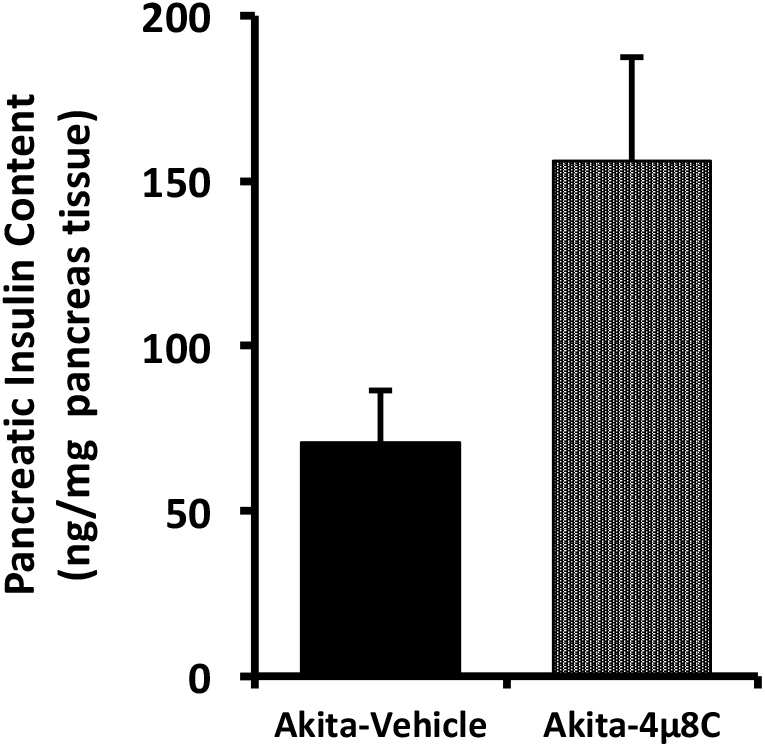
Insulin content measurement of pancreatic tissues treated with vehicle or 4μ8C by ELISA as detailed in Methods and Materials.

**Supplemental Figure 8.**
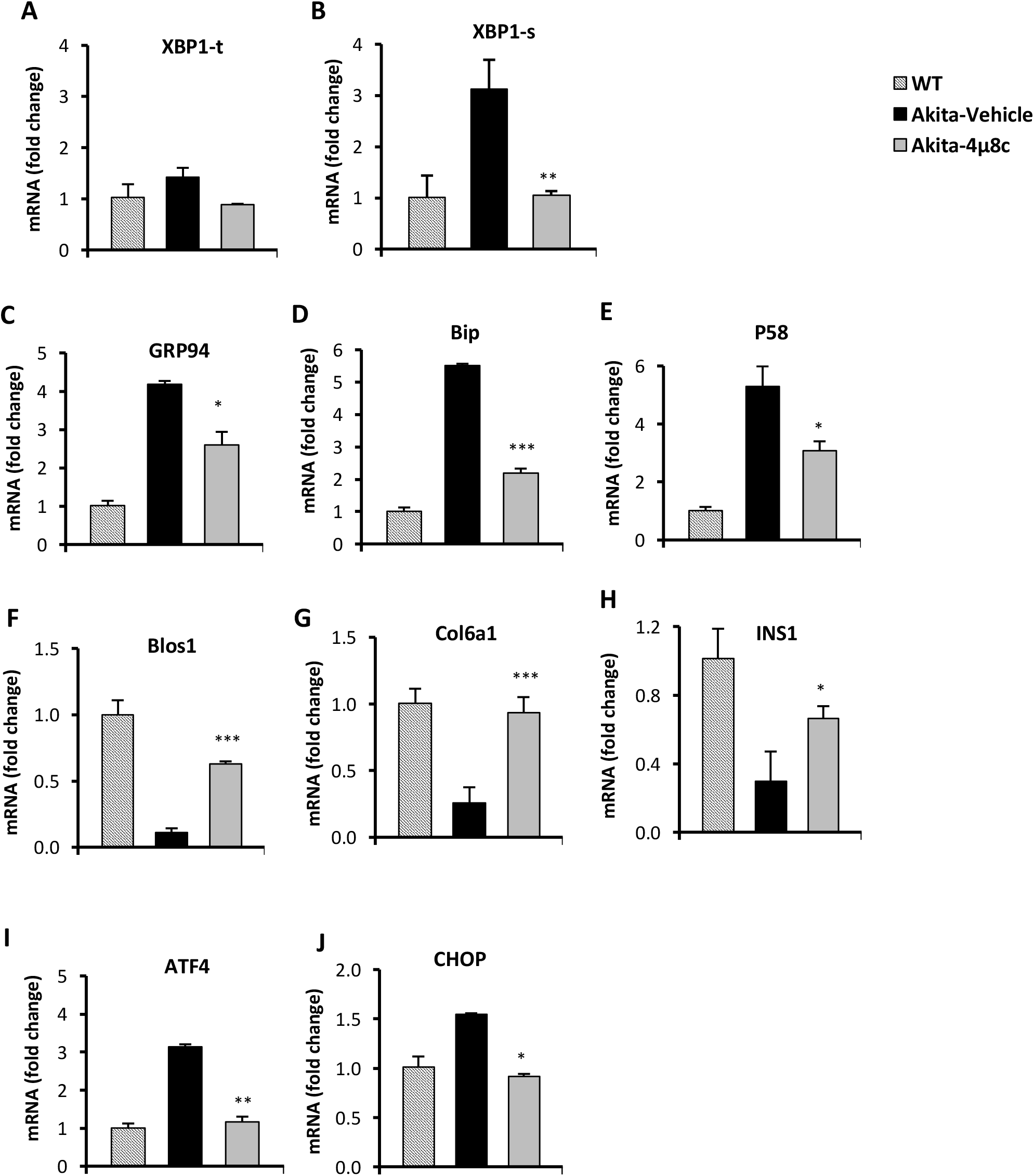
**A-J**. mRNA levels for indicated anti-oxidant genes were analyzed in islets isolated from Akita mice treated with STF or vehicle by qRT-PCR. The results are expressed as fold change and are representative of 3 independent experiments. * P < 0.05, ** P <0.01, and *** P <0.001 compared to Akita-vehicle group. Bars indicate SEM.

**Supplemental Table 1.**
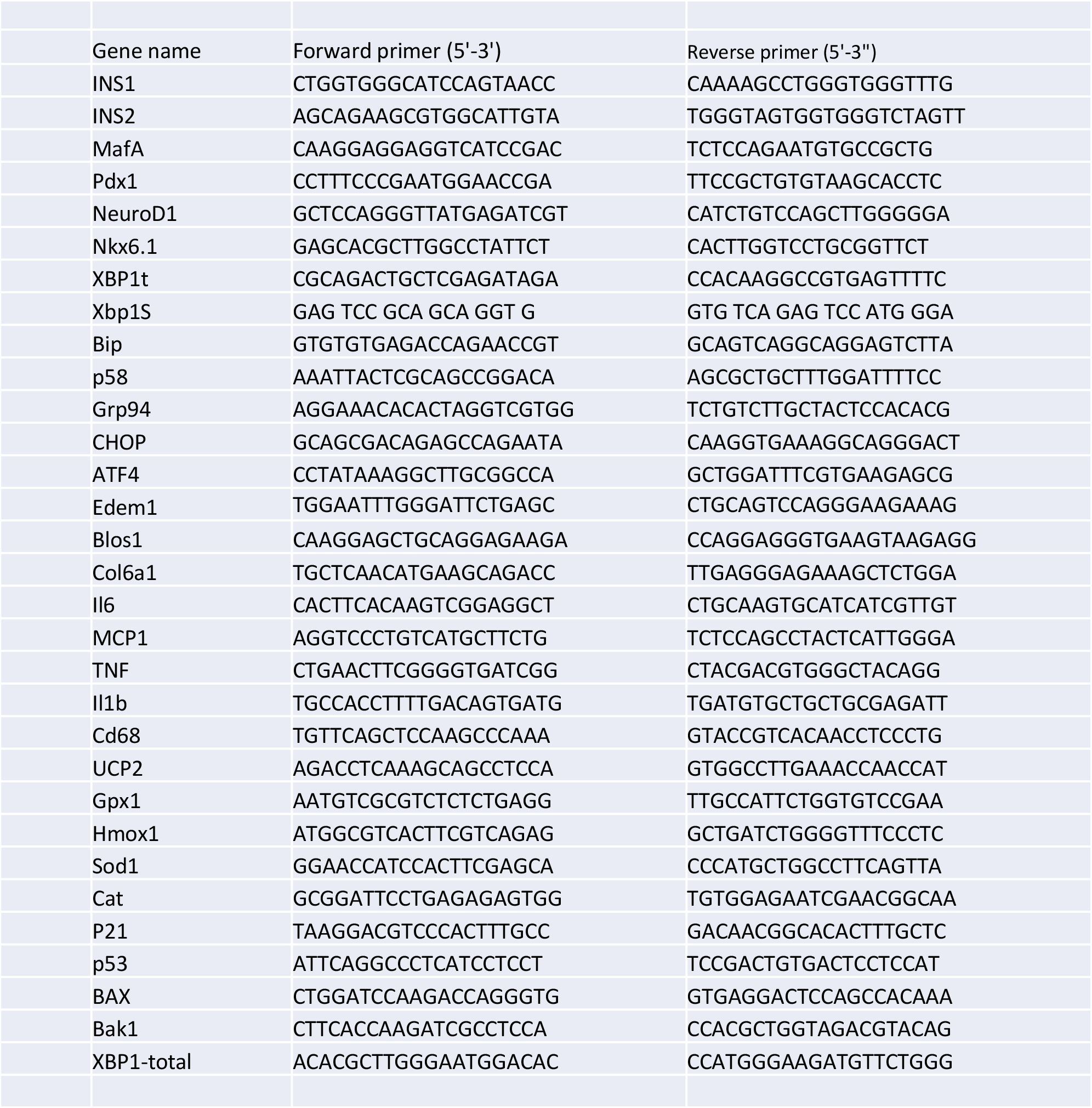

